# Phenology and social status of the endemic Sable Island Sweat Bee, *Lasioglossum sablense*

**DOI:** 10.1101/2023.05.08.539913

**Authors:** Miriam H Richards, Zoe Lucas, Alex Proulx, Lyllian A-J Corbin, Frederica Jacks, Dan Kehler

## Abstract

The Sable Island Sweat Bee (*Lasioglossum (Dialictus) sablense* Gibbs, 2010) is endemic to Sable Island, an isolated sandbar located about 160 km east of Nova Scotia. *L. sablense* is classified as Threatened due to its restricted geographic distribution, so promoting its conservation requires detailed information about nesting phenology and behaviour. We combined measurements and dissections of adult females collected in 2016 and 2017 with behavioural observations of nests and foragers on the grounds of the Sable Island Station in 2019 and 2022, to compile the first description of flight phenology and social status. Like many members of its subgenus, *L. sablense* exhibits a diphasic life history. Phase 1 begins when large adult females emerge from hibernation, begin burrow and brood cell construction, and forage to provision Brood 1, which comprises both daughters and sons. Most, but not all, nests initiated during Phase 1, reactivate during Phase 2, as adult Brood 1 daughters emerge from their nests and initiate foraging to provision Brood 2. This suggests a mix of univoltine (single generation) and bivoltine (two generation) reproductive strategies. Behavioural observations at nest entrances demonstrated that during Phase 2, reactivated nests sometimes contained multiple adult females, suggesting the potential for colonies to become social. Comparisons of body size and ovarian status of Phase 1 and 2 females collected from flowers, showed Phase 2 foragers were significantly smaller than Phase 1 foragers and had somewhat lower levels of ovarian development, as expected if Phase 2 females were workers from eusocial nests. However, five large females collected during Phase 2, which had high levels of wear, likely were Phase 1 foundresses that resumed foraging during Phase 2. Taken together, these observations suggest a mix of phenological and social strategies, including both univoltine and bivoltine life histories, and a mix of solitary and social behaviour among reactivated colonies in Phase 2.

## Introduction

Sable Island sits in the Atlantic Ocean about 160 km east of the Nova Scotia mainland. Approximately 40 km long, with a maximum width of 1.3 km, and comprising a total area of 2,700 ha (∼27 sq km), the island’s dimensions vary as its shoreline is subjected to seasonal and long-term changes in patterns of sand erosion and deposition (Eamer et al. 2022). Geologically, Sable Island is an emergent sandbar that became clear of ice as glaciers retreated after the Last Glacial Maximum, about 12,000 to 13,000 years ago (Stea et al. 1998). Although the island is described as having an impoverished flora and fauna, a number of endemic and geographically restricted plants and animals have been recorded (Howden et al. 1970, Catling et al. 1984, 2009, Mazerolle 2015, Lucas and Brunelle 2016). Five species of bees (Hymenoptera, Anthophila) are found on the island, including two, closely related sweat bee species, *Lasioglossum (Dialictus) sablense* Gibbs, the endemic Sable Island Sweat Bee, and *L. (D*.*) novascotiae* (Mitchell), the Nova Scotia Sweat Bee, which despite its name has a wide range across Canada and eastern North America (Gibbs 2010). In 2014, *L. sablense* was listed as Threatened by the Committee on the Status of Endangered Wildlife in Canada due to its restricted geographic distribution (COSEWIC 2014), and in 2018 it was listed under the Species at Risk Act (SARA; Parks Canada 2020).

In this paper we provide the first study of reproductive phenology and social behaviour of *L. sablense*, combining behavioural observations of bees at their nests in 2019 and 2022, with social trait information (body size and reproductive status) from adult female bees collected prior to COSEWIC and SARA listing. The primary objective of this study was to fill major gaps in knowledge about the species’ reproductive biology, to enhance monitoring and management of this protected species. The second motivation for this study is to provide preliminary behavioural data to be used in comparative studies of social evolution in the subgenus *L. (Dialictus)*, a group in which sociality is clearly an evolutionarily labile trait (Breed 1976, Wcislo et al. 1993, Awde and Richards 2018, Corbin et al. 2021).

## Methods

### Field sites

Sable Island is located 160 km east of the Nova Scotia mainland. Sable Island is an emergent sandbar approximately 40 km long, a maximum width of 1.3 km, and a total area of 2,700 ha (∼27 sq km). There has been a continuous human presence on Sable Island since 1801. Roughly 50% of the island’s land surface is vegetated, and about 185 plant species occur in distinctive vegetation communities comprised of herbs and low shrubs (Colville et al. 2016). Approximately 20% of plant species are introductions, but many are found mostly in areas where human activities and structures provide suitable habitat.

Four locations have been identified as critical nesting habitat for *L. sablense* (Parks Canada 2020, page 17, figure 2). All four are in areas that have for decades been subjected to concentrated human activity within the station compound, including daily foot and vehicle traffic and staging of equipment and cargo. These activities have contributed to soil compaction and the maintenance of sparse vegetation cover at the four sites. The largest of the four nesting sites is in the grounds of the Sable Island Station in the middle of the Island, adjacent to the APU building (Alternate Power Unit), which was built in 1943, and within 10-15 m of a stand of *Rosa rugosa* and is at slightly higher elevation than most other areas of the station (Figure 1). *R. rugosa* appears to be an important forage species for *Lasioglossum*, with as many as ten bees at a time feeding on one flower, especially early in the season (Zoe Lucas, unpub. obs.). Most observations reported here are for nests near the APU building.

**Figure 1.**
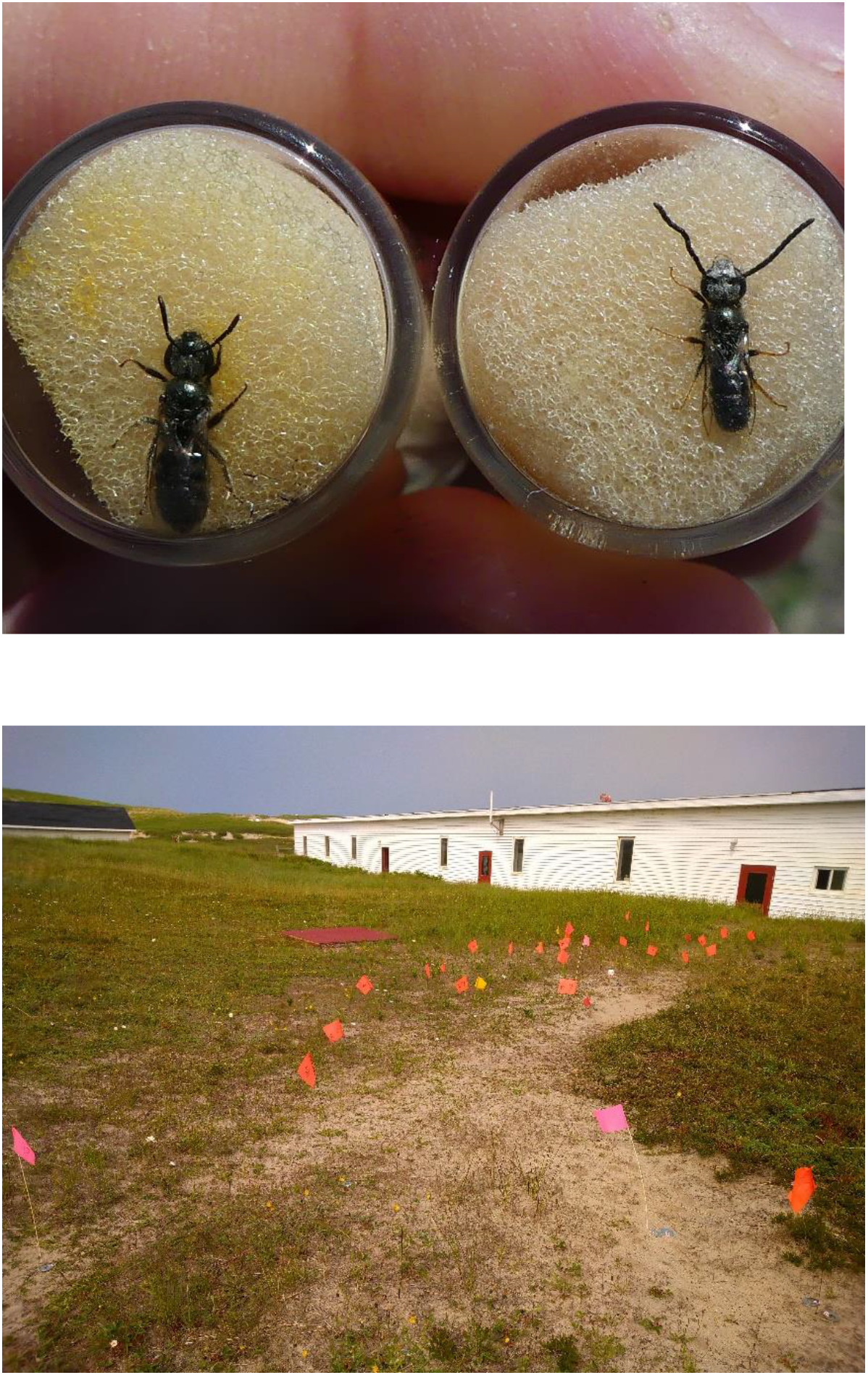
Specimens and nests of *Lasioglossum sablense*, 2019. Top: Female (left) and male (right) specimens of *Lasioglossum sablense* immobilized by sponges in glass vials, allowing for closer examination. Bottom: Nesting aggregation near the APU building at the Parks Canada Station, with nest entrances marked by coloured flags, showing the extent of the aggregation (approximately 8 m). Nests were also marked with flat metal washers, which are not visible in this photo. Note that most nests were found in the well trodden path between buildings.

**Figure 2.**
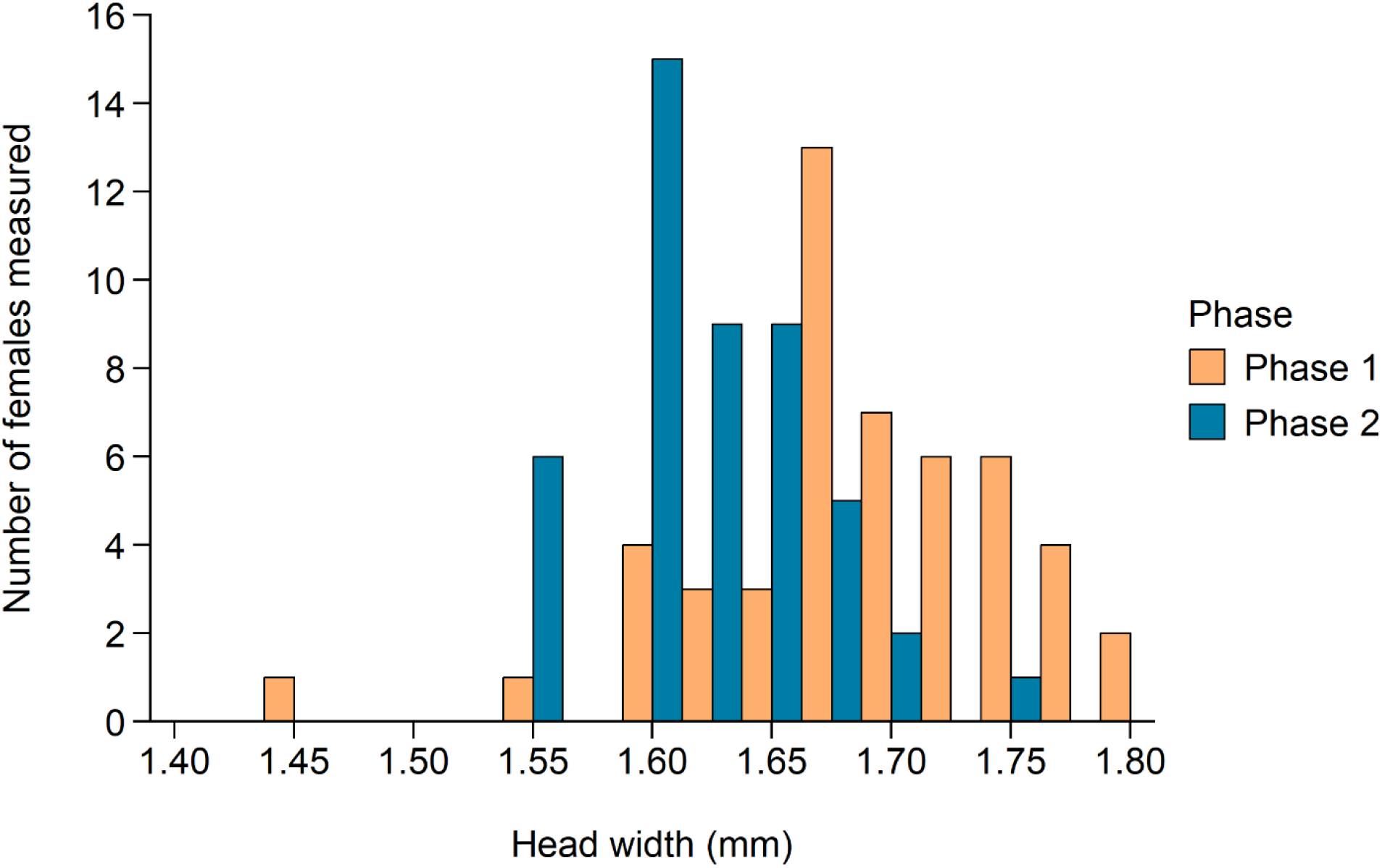
Head width distributions for all available *Lasioglossum sablense* females collected during the first and second phases of the colony cycle in 2016 and 2017. Phase 1 females were significantly larger (one-way ANOVA, adj. R^2^=0.26, F=33.83, df=1,95, p=8.08e-08).

### Specimen collections

All *L. sablense* specimens used for dissections were collected prior to recognition of the species’ status under SARA in 2018. Two *Lasioglossum sablense* specimens were collected near the APU building at the Sable Island Station on 1 and 2 June 2016, using a yellow bowl trap filled with water and a drop of dishwashing detergent, placed on the ground?. Another 97 specimens were collected by targeted netting during surveys of forage plant species for *Lasioglossum* (both *sablense* and *novascotiae*) on 2 June 2016, 2 to 4 August 2016, and 1 to 6 August 2017. Plant species surveyed were *Vaccinium macrocarpon* (Ericaceae), *Rosa virginiana* (Rosaceae), *Sibbaldiopsis tridentata* (Rosaceae), *Achillea millefolium* (Asteraceae), *Leontodon autumnalis* (Asteraceae), and *Leucanthemum vulgare* (Asteraceae), at seven locations from the Sable Island Station (43.9329N, −60.0074W) eastward to 43.9483N, −59.8187W. Bees foraging on flowers were collected into vials of 70% EtOH and within 24 hours were placed in glass vials with fresh 70% EtOH for storage, then sent to the Nova Scotia Museum of Natural History for identification. *Lasioglossum sablense* and *L. novascotiae* can be reliably distinguished from each other by the density of punctures on the mesoscutum (Gibbs 2010).

### Observations of bees at nests

An aggregation comprising at least 95 nests was discovered along a footpath between the APU building and the garage at the Sable Island Station on 3 July 2019 (diagram of site). The nesting aggregation covered a sandy area roughly 2 m wide and 10 m long, and most nests were found along the edges of the packed footpath. From 3 to 9 July, nests were identified when a female was observed departing from or arriving at the entrance. Additional observations were carried out at nests near the APU building, between 8 and 15 August 2022, using similar methods.

Nest entrances were marked with short flags or with flat steel washers to which a piece of tape was attached with the nest number (Figure 1). The washers were usually set down after a forager entered the nest, to avoid confusing females, which often did orientation flights around the nest entrance on their next foraging flight. Although some nests had temporary tumuli, after nest entrances were closed at the end of the day, the tumuli were easily obliterated by wind or rain, so the number of nest entrances was likely underestimated. Occasionally, two nest entrances were found within a few millimetres of each other, and these were assumed to represent separate nests, rather than one nest with two entrances.

To identify which of the two *Lasioglossum* species were the subject of our observations in 2019, we caught 10 females nesting across the main aggregation and examined them briefly with a 10x eyepiece micrometer before releasing them; all 10 were identified as *L. sablense*. In 2019, we found females of *L. novascotiae* nesting in the vicinity, in small patches of sandy soil near other buildings 40 to 100 m away from the main aggregation, but not in the main sweat bee nesting aggregation near the APU or in three other small nesting aggregations in the vicinity. In 2022, bees from six nests were caught and individually marked for observations at nest entrances; two were identified as *L. sablense* and four as *L. novascotiae*, suggesting that by 2022, the APU aggregation was comprised of nests of both species.

### Phenology and behaviour at nest entrances

Female bees were observed at nest entrances during short observation periods (30 min to 4 hours) on 3 to 10 July, 22 July, and 5 to 9 August in 2019, and on 8, 10, 12, 14 and 15 August in 2022. Observations were aimed mainly at determining whether single or multiple females were resident or foraging, so in 2022, a few adult females were caught and individually marked on the thorax with enamel paint in order to facilitate behavioural observations at nest entrances.

We recorded behaviours typically collected in nest-based studies of sweat bee sociality: whether nest entrances were open or closed, departures of adult females from their nests (including orientation flights), arrivals of adult females at nest entrances with or without pollen loads, searching behaviour by adult females, and nest digging. In 2019 we also carried out short visual surveys (15 to 30 minutes) of sweat bees foraging on flowers growing within the station precincts on 27 June, 16 July, 22 July, and 9 August. In 2022, we netted bees foraging on flowers within the station precinct, on 6 days from 8 to 15 August, in order to briefly examine them and to measure head width, wing wear, and mandibular wear, before release.

### Social status of *L. sablense*

A total of 97 female specimens was collected in 2016 and 2017. For these 97 specimens, we measured head width, mandibular wear and wing wear, as these can be measured without damaging specimens. As *L. sablense* specimens can no longer be collected, we dissected only a subset of 30 specimens, including three whose heads were detached, prioritizing females with higher wing wear scores, as these were most likely to have been engaged in brood provisioning activities. Measurement and dissection methods follow Corbin et al. (2021). Briefly, head width (HW) was measured as the widest distance across the head, including the compound eyes, in millimetres. To measure costal vein length (CVL), we detached the wings, taped them to paper with clear tape to hold them flat, and then measured the length of the costal vein in millimetres. Wing wear and mandibular wear were each measured on a scale from 0 (no wear) to 5 (very worn), and then summed to create a total wear score (TW) for each specimen. Females were dissected and their ovarian status was evaluated in two ways. First, the sizes of all developing oocytes relative to a fully developed egg ready to lay (1/4, 1/2, 3/4, and 1) were summed to create a volumetric ovarian development score (OD). Second, we scored females according to the fractional size of their largest developing oocyte; those with at least one 1/2-developed oocyte were classified as ‘fecund’ (Breed 1976). The heads of three specimens were detached during transport so could not be positively associated with the dissected abdomens.

## Results

### Phenology

Observations at nest entrances in 2019 suggest two successive periods of brood provisioning activity. From 5 to 7 July 2019, more than half of nest entrances were open at least once during the day (Table 1). By 9 July and again on 22 July, fewer than 15% of marked nests were open, suggesting that the first brood-provisioning phase had finished, with foundresses remaining in their burrows until the emergence of their first workers. Throughout the month of July, we only ever observed a single female in each nest. In early July, we also observed females engaged in the low, sinuous flight patterns characteristic of searching behaviour, in which females fly back and forth close to the ground, occasionally landing and crawling about on the sand and under vegetation, inspecting the surface.

**Table 1.** Proportions of *Lasioglossum sablense* nests that were active during occasional surveys in 2019. Nests were recorded as active if the nest entrance was open or if a female was observed departing or arriving with pollen. The decline in activity from 9 to 22 July suggests a quiescent period between B1 and B2 provisioning phases.

| Date | Nest entrances<br>observed | Nests with<br>activity (%) |
| --- | --- | --- |
| 5 July | 67 | 50 (75) |
| 6 July | 24 | 16 (67) |
| 7 July | 20 | 11 (55) |
| 9 July | 21 | 3 (12) |
| 22 July | 68 | 10 (15) |
| 5 August | 57 | 26 (46) |

These females likely were nest foundresses looking for suitable nest excavation sites. Supporting this interpretation, there were several instances in which we marked and observed a single nest entrance, but subsequently found two separate nest entrances.

On 5 August 2019, almost half of nests reopened (Table 1), and foragers were again observed departing and arriving at nest entrances. In August 2019, some nests contained multiple females; for instance, at least three females entered nest 63 within the span of one minute on 5 August. Visual surveys of bees on flowers around the Field Station on 27 June, 16 July, and 22 July 2019 indicated that all foragers were females, whereas on 9 August that year, males were also observed foraging on flowers; since male halictids do not overwinter, these must have been newly eclosed. Similarly, observations and netting of specimens from 8 to 15 August 2022 yielded both females and males foraging on flowers.

One male examined on 11 August 2022, had highly worn wings (WW=5), suggesting considerable flight activity, whereas three other males caught the same day showed no wing wear (WW=0), and likely were newly eclosed. These observations suggest that nests were reactivated in early August following eclosion of Brood 1. In a mix of marked *L. sablense* and *L. novascotiae* nests surveyed in August 2022, about 70% had either nest guards or foragers. More detailed observations of four active nests definitely known to belong to *L. sablense* between 8 and 15 August 2022, demonstrated that multiple females may be active simultaneously. In one nest, for instance, at least three unmarked foragers arrived at the nest in quick succession, one of them carrying pollen. Based on observations of these four nests, there seemed to be no more than one female carrying pollen in each nest, and the maximum number of pollen trips observed per day was only three.

### Female traits related to sociality

Based on the bivoltine phenology inferred above, we assumed that adult females observed or collected in June or July (Phase 1) were nest foundresses, while those collected or measured in August (Phase 2) were B1 daughters. Phase 1 females were significantly larger than those collected in Phase 2 (Figure 2). Based on the difference in average head width, Phase 2 females were 1.8% smaller than Phase 1 females (Table 2).

**Table 2.** Trait comparison for *Lasioglossum sablense* females collected during Phases 1 and 2 in 2016 and 2017. The proportional size difference was calculated as (Phase 1 – Phase 2) / Phase 1 × 100. Head width and costal vein length are given in mm.

| Trait | Phase 1 | Phase 2 |
| --- | --- | --- |
| Head width (mean ± s.d.) | 1.68 ± 0.05 | 1.65 ± 0.06 |
| Costal vein length (mean ± s.d.) | 1.90 ± 0.06 | 1.88 ± 0.08 |
| Total wear score (median, range) | 3 (1 – 6) | 4 (2 – 10) |
| OD score (mean ± s.d.) | 0.98 ± 0.67 | 0.44 ± 0.46 |
| Proportion fecund | 14 / 15 = 93% | 11 / 15 = 73% |

We analysed relationships between size (HW) and wear of foraging females collected in Phases 1 and 2. Total wear scores (TW) were higher during Phase 2 (Table 2). Moreover, TW scores also seemed to be higher in larger females (Figure 3, Table 3). However, this relationship appeared to be driven by the very high wear scores of the five largest females collected during Phase 2, which likely were Phase 1 nest foundresses that began a second bout of foraging during Phase 2. When these were excluded from consideration, the significant association between size and wear disappeared, as did the difference in total wear between Phases 1 and 2 (Table 3).

**Table 3.**
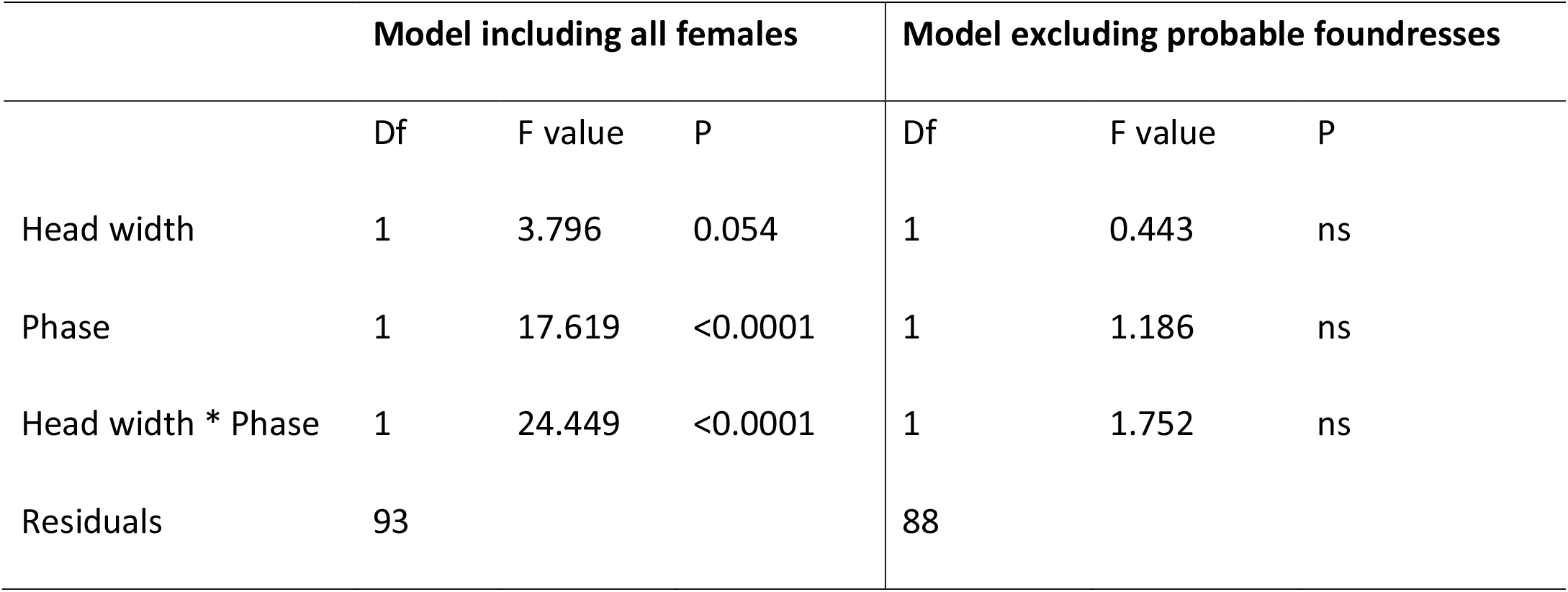
Linear models analysing the relationship between head width and total wear scores in Phase 1 and Phase 2 females (see Figure 2). The association between size and wear was significant, especially during Phase 2, when all females were included, but not when the five probable foundresses collected in Phase 2 were excluded.

**Figure 3.**
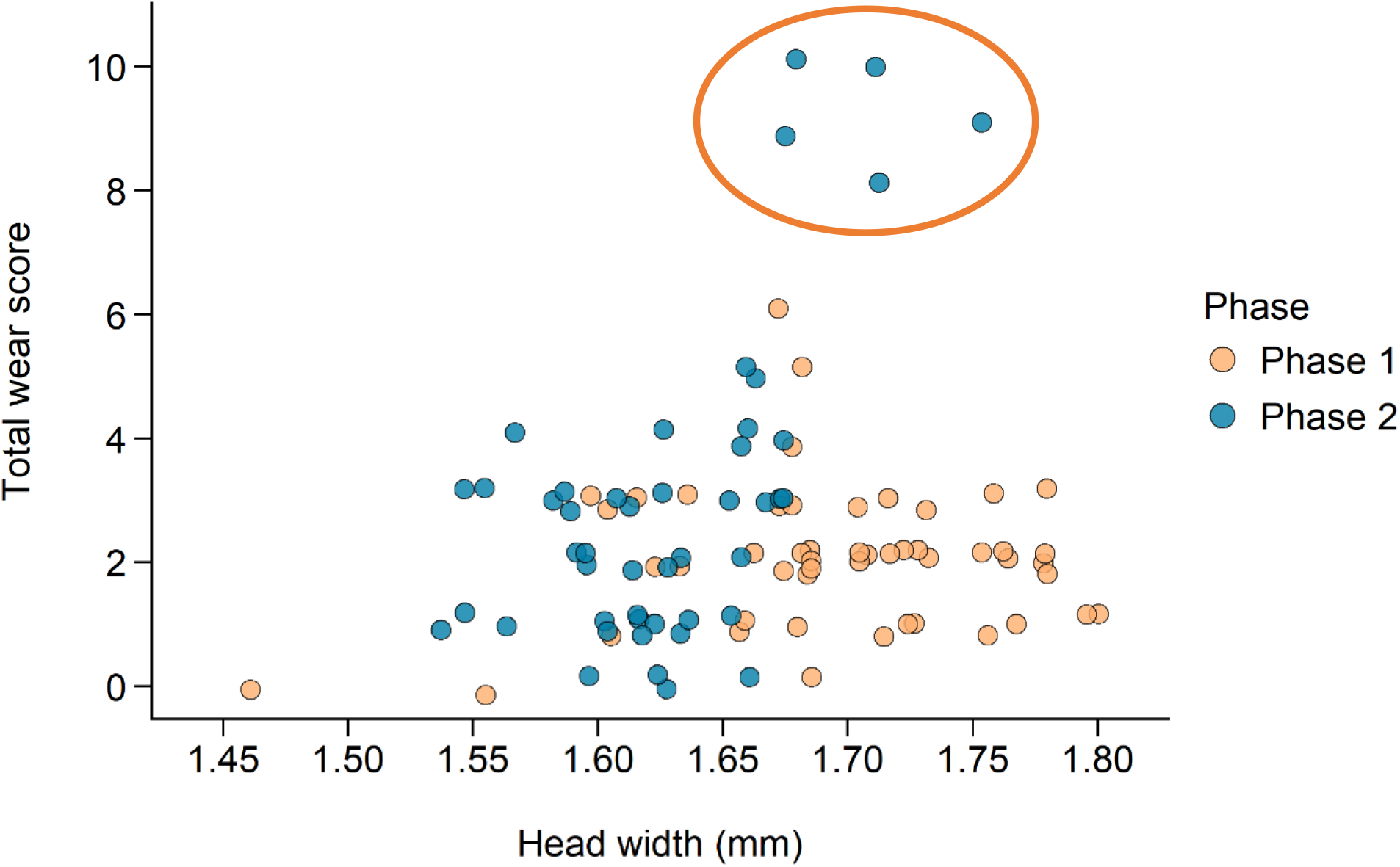
Relationship between body size and wear in *L. sablense* females. Phase 2, but not Phase 1 females, showed a significant association between size (HW) and wear (TW), driven by five highly worn females collected in Phase 2 and indicated with a circle.

Phase 2 females tended to have lower ovarian scores than Phase 1 females (Figure 4), although the proportions of fecund females (with largest oocyte at least 1/2-developed) were similar in Phases 1 and 2 (Table 2, Fisher exact probability = 0.33). Since Phase 1 females tended to be larger and have higher ovarian scores than Phase 2 females, we also analysed relationships between size and ovarian development using a linear model of the form OD ∼ size + phase. Ovarian score was not significantly associated with head width (adj. R^2^=0.09, F=2.43, df=2,27, n.s.), but was significantly associated with costal vein length (adj. R^2^=0.15, F=3.48, df=2,27, p=0.045; Figure 5).

**Figure 4.**
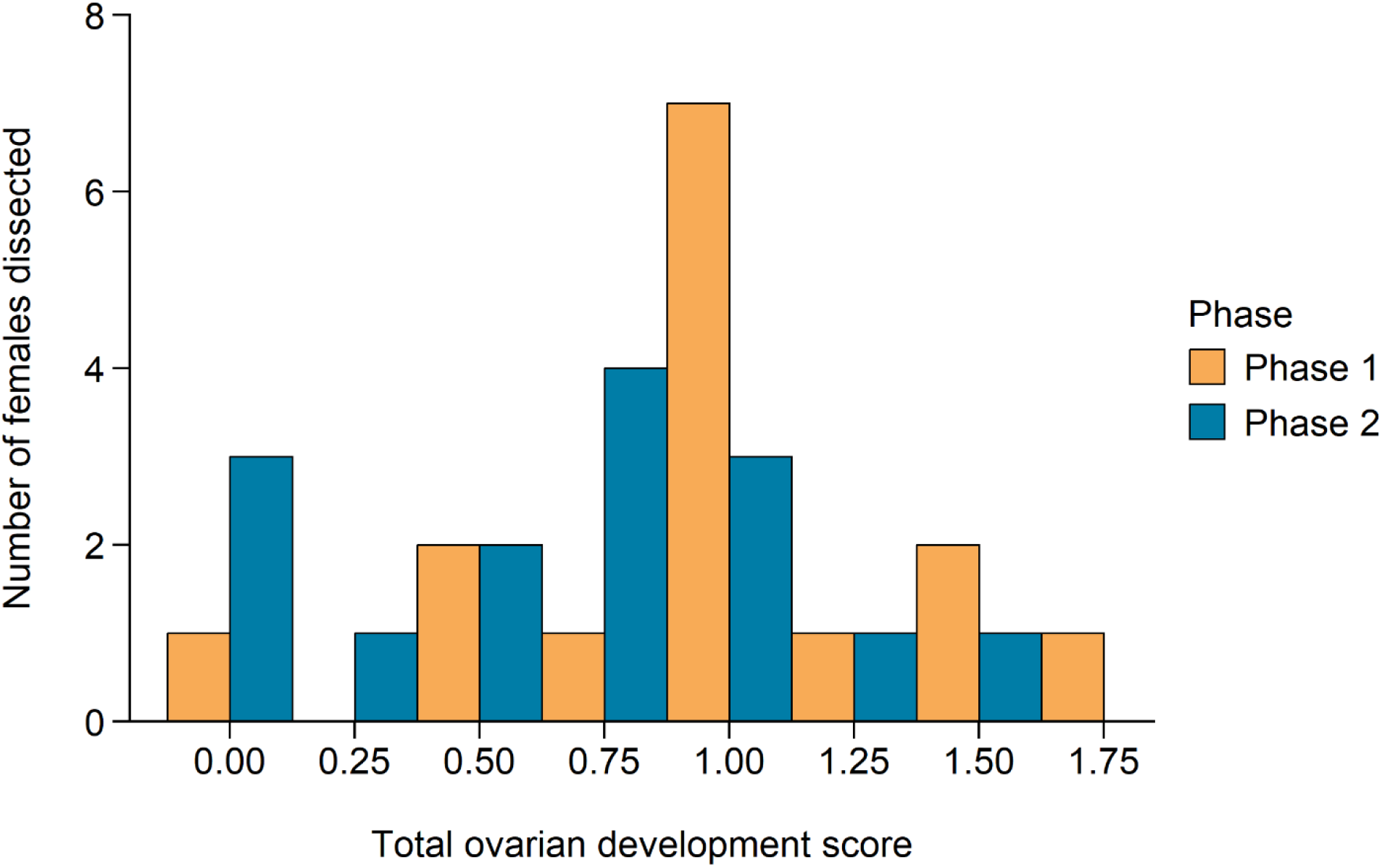
Comparison of ovarian development scores for *Lasioglossum sablense* females collected in 2016 and 2017. On average, phase 1 females tended to have higher OD scores than Phase 2 females (one-way ANOVA, adj. R^2^=0.09, F=4.04, df=1,28, p=0.054).

**Figure 5.**
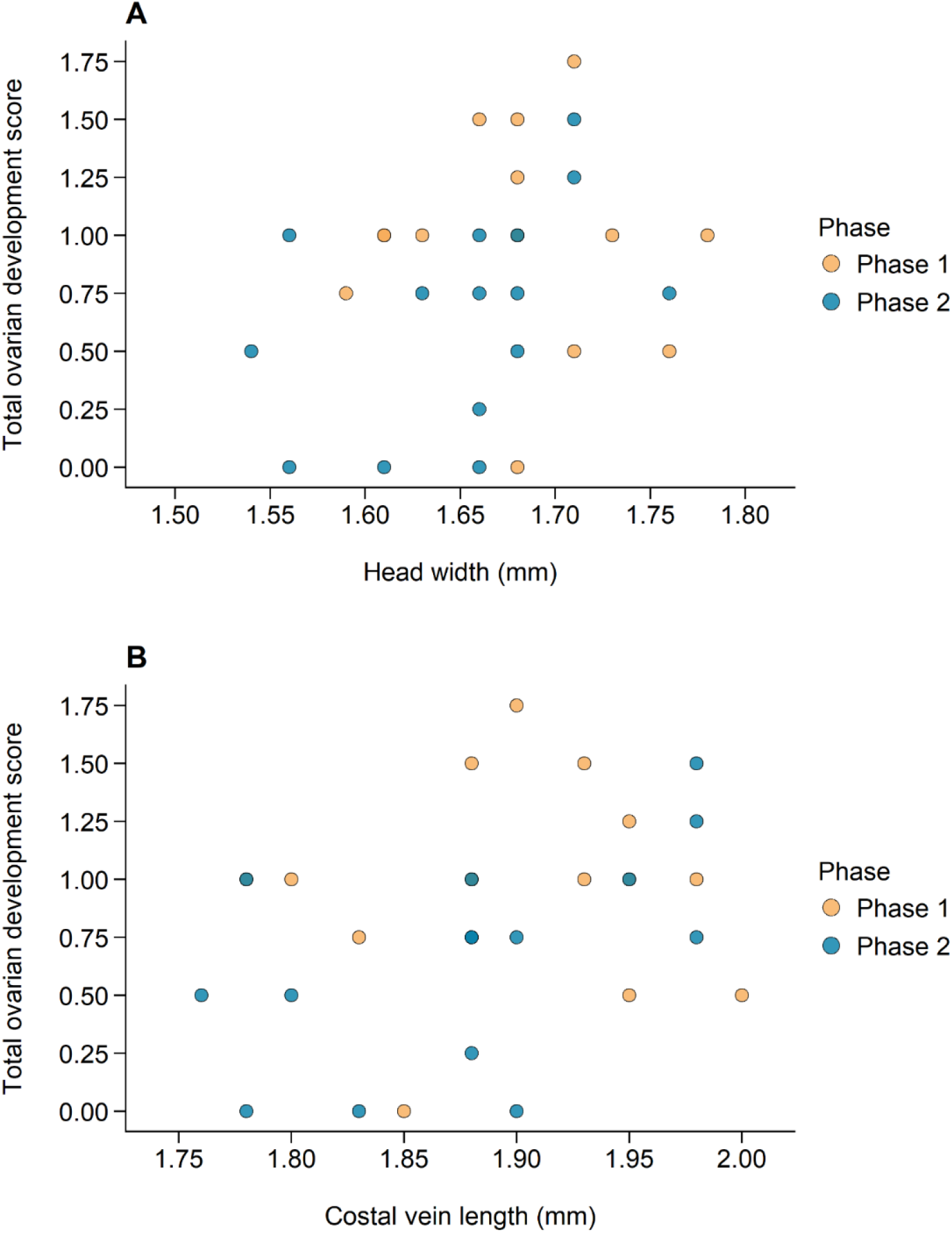
Variation in OD scores based on body size and phenology for female *L. sablense* collected in 2016 and 2017. A. Size measured by head width. B. Size measured by costal vein length.

In eusocial sweat bees, altruistic workers can be recognized by a combination of high wear and no ovarian development. There were four females with OD=0 and TW>1 among the dissected specimens, one collected in Phase 1 (with TW=6) and three collected in Phase 2 (with TW=2, 4, and 4).

## Discussion

### Phenology and social status of *Lasioglossum sablense*

Based on the preliminary studies presented here, we suggest the following summary of *L. sablense* phenology and social behaviour. Foundresses begin to establish nests solitarily in Phase 1, likely in June. Nest founding is asynchronous, and new nests may be initiated as late as early July. Most foundresses finish provisioning Brood 1 (B1) by early July, and close their nest entrances, remaining inside the nests until the emergence of B1 adults in early August. Brood 1 comprises both females and males, and their emergence as adults marks the beginning of Phase 2 of the colony cycle. During Phase 2, several categories of adult females are active outside their nests. Most of these are Brood 1 daughters that remain in the maternal nest, but a few are old foundresses that resume foraging to provision a second brood.

The activity of adult females during Phase 2 is key to understanding the nature of colony social organisation in bivoltine sweat bees (Schwarz et al. 2007, Corbin et al. 2021). We observed several nests in which multiple females were active outside the nest, but no nests in which there were multiple females collecting pollen provisions simultaneously. Given that only a few nests that we observed in 2022 were definitely occupied by *L. sablense*, it would be premature to hazard a guess at the proportion of Brood 1 daughters that become foragers during Phase 2. Nevertheless, our observations do demonstrate that most nests reactivate during Phase 2, and that in many of these, Brood 1 daughters become foragers. The available evidence suggests several different types of social behaviour occur in *L. sablense*. First, some colonies seem to be functionally solitary. Even in obligately eusocial sweat bees, functionally solitary behaviour occurs when no Brood 1 females survive to adulthood, either ending the colony cycle during Phase 1, or forcing nest foundresses to resume brood provisioning during Phase 2 (Richards and Packer 1994). Colonies may also be considered as functionally solitary if only a single daughter provisions Brood 2 in the maternal nest, while other daughters either remain leave or enter hibernation.

Alternatively, colonies may become multi-female associations during Phase 2. Most Phase 2 foragers had developing ovaries, suggesting they were provisioning their own eggs, amd that multifemale associations are often communal. However, in our sample of dissected Phase 2 females, we found some small worn females with no ovarian development, a trait combination suggesting these were workers. Since foundresses evidently can live long enough to forage during Phase 2, this implies the potential for colonies to become eusocial, with daughters provisioning brood that develop from foundress-laid eggs. If so, then *L. sablense* should be regarded as ‘weakly eusocial’, a term that has been used to describe obligately eusocial sweat bees in which workers exhibit high levels of ovarian development, as in an Alberta population of *L. laevissimum* (Packer 1992). The proportion of fecund B1 daughters in *L. sablense* (84%) is the highest of any *Dialictus* yet studied. In some ways, the behaviour of *L. sablense* resembles that of the primarily solitary species, *L. figueresi* Wcislo (Wcislo et al. 1993), in which sociality occurs when B1 daughters remain together in their natal nest. Usually, each B1 daughter provisions her own brood and the colony becomes communal, but very rarely, one female lays eggs that are provisioned by her sisters, creating a semisocial colony. *L. figueresi* foundresses almost always die before their daughters emerge, so colonies cannot become eusocial, whereas in *L. sablense*, the long lives of foundresses do create the potential for eusociality.

## Conclusions

The preliminary data presented here provide a tantalizing outline of the phenology and social behaviour of *L. sablense*. The Sable Island Sweat Bee is part of the *viridatum* group of *Dialictus* (Gibbs 2010), which includes the only other sweat bee found on Sable Island, the behaviourally uncharacterized *L. novascotiae*. Related species include some well studied eusocial species such as *Lasioglossum laevissimum* (Packer 1992, Awde and Richards 2018), which seem to be obligately eusocial and thus less socially variable than *L. sablense*. More detailed studies are required to evaluate the frequencies of alternative social strategies in this species, to test the hypothesis that it is, in fact, socially polymorphic, exhibiting an array of social strategies, including solitary, communal, eusocial, and possibly semisocial behaviour. The fact that *L. sablense* is endemic to an isolated island that is only about 10,000 years old, suggests that it evolved as a separate species quite quickly and so might become a fascinating model for investigating the time course of social evolutionary change in halictine bees.

## Acknowledgments

We thank John Klymko for help finding bee nests around the Sable Island Station in 2019, Dominique Gusset, Sue Westby, and Jason Gibbs for species identifications, the Nova Scotia Museum of Natural History, Halifax, Nova Scotia for allowing us to dissect specimens. Funding and logistical support were provided by Parks Canada, the Sable Island Institute, and the Natural Sciences and Engineering Council of Canada.

## Notes

### Competing Interest Statement

The authors have declared no competing interest.

